# Immunochemical analysis of the integrin expression in retinal pigment epithelial cells

**DOI:** 10.1101/2020.06.26.133553

**Authors:** Janosch P Heller, Tristan G Heintz, Jessica CF Kwok, Keith R Martin, James W Fawcett

## Abstract

Retinal pigment epithelial (RPE) cells have been used in disease modelling and transplantation studies in retinal diseases. Several types of RPE cells have been trialed, ranging from primary cells and immortalized cell lines to stem cell-derived RPE cells. During aging and in disease, the extracellular environment of the RPE cells changes, interfering with RPE cell adhesion. We hypothesize that this could be a key problem in transplantation studies that have reported lack of adhesion and survival of the transplanted RPE cells. Integrins are essential for the proper function of the RPE, mediating adhesion to Bruch’s membrane, and the binding and subsequent phagocytosis of photoreceptor outer segments. Variability that has been found in clinical trials might be due to the variability of cell types used and their expression profiles of surface molecules. Here, we set out to analyze integrin expression in primary rat RPE cells and in the human cell line ARPE-19 using immunochemistry. We found that both cell types express integrins to varying degrees. After long-term culturing, ARPE-19 cells resemble mature RPE cells, and increase integrin expression. We hence argue that it is important to test the properties of these cells prior to transplantation to avoid failure of adhesion and to facilitate correct function.

## Introduction

The retinal pigment epithelium (RPE) is located behind the neural retina, forming a barrier between the photoreceptor cells and the blood vessels of the choroid. RPE cells are critical for the development and integrity of the outer retina. They transport nutrients from blood vessels to photoreceptors and phagocytose shed photoreceptor outer segments (POS) as well as playing a role in maintaining the blood-retina barrier [1–3]. Integrins are crucial for the proper functioning of the RPE cells. The adhesion of the RPE monolayer to its normal substrate, Bruch’s membrane, is mediated via integrins [4,5]. Additionally, αVβ5 integrin is essential for POS binding and subsequent phagocytosis through CD36 and the proto-oncogene tyrosine-protein kinase MER (MERTK) [3,6–9]. Integrins are transmembrane signaling molecules that are formed through the hetero-dimerization of an α and a β subunit [10,11]. They are important adhesion molecules, binding to a variety of extracellular matrix (ECM) cues as well as cell surface markers and growth factors [12,13].

During aging and in disease state, the ECM composition of Bruch’s membrane changes and, together with accumulation of toxic waste due to a decline in the efficiency of phagocytosis and subsequent recycling and degradation may lead to the dysfunction and ultimate death of RPE cells [14–17].

RPE cell transplantation has been employed as a treatment to replace damaged or lost RPE cells in retinal diseases such as Stargardt disease, age-related macular degeneration (AMD) and retinitis pigmentosa. However, attempts to transplant new RPE cells into diseased eyes of human AMD patients have been proven challenging [18–25]. Even though the transplantation resulted in improved vision in some cases, most of the grafts did not attach and survive in the pathological microenvironment due to immune rejection and failed adhesion [19,26–32]. Early results from the recent clinical trials involving stem cell-derived RPE cells are promising as no signs of adverse effects such as tumorigenesis were described [33–35]. Despite the reported improved vision in patients it is still unknown whether the transplanted cells adhere properly to the pathological Bruch’s membrane and whether they form a polarized monolayer *in vivo* [36].

Several different types of RPE cells have been used to study the behavior of the cells *ex vivo* prior to transplantation or to model diseases. Cell lines [37–40], including the widely used human RPE cell line ARPE-19 [41], and primary RPE cells such as fetal [42,43] and adult human cells [44] as well as stem cell-derived RPE cells [45–48] were used in transplantation experiments in animal models with varying degrees of success [49].

One of the problems with the *ex vivo* expansion of RPE cells prior to transplantation is dedifferentiation of the cells. For example, ARPE-19 cells express RPE-specific genes to a reduced degree [50] but upregulate neuronal markers [51]. In addition, the dissociation of RPE cells prior to transplantation can lead to the dedifferentiation of the cells and they then have to redifferentiate *in vivo* [46,47,52–54].

Here, we assessed the integrin expression in rat RPE cells *in situ* and in culture as well as in differentiated and undifferentiated ARPE-19 cells by immunocytochemistry to get a better understanding of the abilities of the different variety of RPE cells to attach to Bruch’s membrane after transplantation.

## 2. Material and Methods

### 2.1 Cell culture

#### 2.1.1 Primary adult rat RPE cells

All animal work was carried out in accordance with the UK Animals (Scientific Procedures) Act 139 (1986) and within UK Home Office regulations. Adult Sprague Dawley (SD) rats (Charles River; 250–300 g, 8–10 weeks) were used for tissue. Primary adult rat RPE cell cultures were prepared as described previously [55]. In brief, whole eyes were incubated in papain (Worthington; 20 U/ml) for 50 minutes at 37°C. Afterwards, the retina with attached RPE cells was dissected out, and the rest of the eye was discarded. After a second digestion of the retina-RPE complex in papain (20 U/ml) for ten minutes at 37°C the RPE cells were carefully peeled off and plated on Matrigel (BD Biosciences; 1:80 in DMEM)-coated glass coverslips in ‘Miller’ medium [DMEM supplemented with 5 or 20% fetal bovine serum (v/v), N1 medium supplement, MEM non-essential amino acids, 2 mM GlutaMAX, 250 mg/ml taurine, 20 ng/ml hydrocortisone, 13 ng/ml triiodothyronine and 1% penicillin/streptomycin (v/v); 56,57].

#### 2.1.2 ARPE-19 cells

ARPE-19 cells [41] were purchased from ATCC. Cells were plated on cell culture-treated plastic in DMEM^+^ [DMEM, high glucose, pyruvate, supplemented with 1% fetal calf serum (v/v) and 1% penicillin/streptomycin (v/v); 58]. Cells were fed twice a week and split when confluent. When kept in culture without splitting cells became pigmented after ~five months.

### 2.2 Immunochemistry

#### 2.2.1 Immunocytochemistry

ARPE-19 cells were briefly trypsinized and plated in small number on Matrigel-coated glass coverslips. We used undifferentiated (cultured for two weeks) and differentiated (cultured for ~five months) ARPE-19 cells as well as two week old primary rat RPE cell cultures for the immunostaining.

Cells were fixed with either 2% (integrin staining) or 4% (RPE marker staining) paraformaldehyde (PFA) for 15 min at room temperature and washed three times with PBS. Subsequently, the cells were permeabilized and blocked using PBS supplemented with 0.3% Triton-X 100 (PBST) and 5% milk (integrin staining) or 10% donkey serum (RPE marker staining) for one hour. Afterwards, cells were incubated in primary antibody solution (antibody diluted in 5% milk or 5% donkey serum in PBST (Table 1)) over night (~14 to 16 hours) at 4°C. The next day, cells were incubated in secondary antibody solution (antibody diluted in PBST (Table 2)) for one hour. Afterwards, the cells were washed and mounted onto slides for imaging.

**Table 1:** Primary antibodies used. All antibodies are IgG. RRID = Research Resource Identifier.

| Antigen | Host | clone | Supplier | product code | RRID |
| --- | --- | --- | --- | --- | --- |
| BEST | mouse | monoclonal | Millipore | MAB5466 | AB_2064603 |
| CRALBP | mouse | B2 | Novus | NB100-74392 | AB_1048601 |
| Cytokeratin 18 | mouse | C-04 | Abcam | ab668 | AB_305647 |
| integrin α1 | rabbit | polyclonal | Millipore | AB1934 | AB_91199 |
| integrin α2 | rabbit | polyclonal | Millipore | AB1936 | AB_2265143 |
| integrin α3 | rabbit | polyclonal | Millipore | AB1920 | AB_11213115 |
| integrin α4 | rabbit | polyclonal | Millipore | AB1924 | AB_91181 |
| integrin α5 | rabbit | polyclonal | Millipore | AB1928 | AB_2128185 |
| integrin α6 | rat | GoH3 | BD Pharmingen | 555734 | AB_2296273 |
| integrin α8 | rabbit | polyclonal | Santa Cruz | sc-25713 | AB_2265044 |
| integrin αV | rabbit | polyclonal | Millipore | AB1930 | AB_11213475 |
| integrin β1 | rabbit | polyclonal | Millipore | AB1952 | AB_91150 |
| integrin β3 | rabbit | polyclonal | Millipore | AB2984 | AB_10806204 |
| integrin β4 | rabbit | polyclonal | Millipore | AB1922 | AB_91186 |
| integrin β5 | rabbit | polyclonal | Millipore | AB1926 | AB_91119 |
| integrin αVβ3 | mouse | 27.1 (VNR-1) | abcam | ab78289 | AB_1566022 |
| integrin αVβ5 | mouse | P1F6 | abcam | ab24694 | AB_448231 |
| MiTF | mouse | C5 | abcam | ab12039 | AB_298801 |
| Na<sup>+</sup>/K<sup>+</sup>-ATPase | mouse | C464.8 | Millipore | 05-382 | AB_309708 |
| RPE65 | mouse | 401.8B11.3D9 | Millipore | MAB5428 | AB_571111 |
| ZO-1 | rabbit | polyclonal | Thermo Fisher | 61-7300 | AB_138452 |

**Table 2:** Secondary antibodies used. All antibodies are polyclonal and were purchased from Thermo Fisher. RRID = Research Resource Identifier.

| Antigen | Feature | Host | product code | RRID |
| --- | --- | --- | --- | --- |
| mouse IgG | Alexa488-fluorochrome conjugated | donkey | A-21202 | AB_141607 |
| mouse IgG | Alexa568-fluorochrome conjugated | donkey | A-10037 | AB_2534013 |
| rabbit IgG | Alexa488-fluorochrome conjugated | donkey | A-21206 | AB_141708 |
| rabbit IgG | Alexa568-fluorochrome conjugated | donkey | A-10042 | AB_2534017 |
| rat IgG | Alexa488-fluorochrome conjugated | goat | A-11006 | AB_2534074 |

#### 2.2.2 Immunohistochemistry

Adult SD rats (Charles River; 250–300 g, 8–10 weeks) were perfused transcardially with 4% PFA in 0.1 M PBS, pH 7.2-7.4, after overdose of general anesthetic (Euthatal (200 mg/ml solution, Rhône-Mérieux)). Afterwards, eyes were dissected out and excess connective tissue and muscle attachments were removed. The lens and anterior chamber were carefully removed together with the vitreous humor. Care has to be taken to not destroy or detach the retina. The eyes were immediately post-fixed in 4% PFA overnight at 4°C. The tissue was transferred into 30% sucrose (Sigma) in PBS, for cryoprotection. The eyes were then frozen in O.C.T. (Raymond Lamb) and sectioned using a cryostat (Cryostat Leica CM 3050S). Fourteen μm thick sections were mounted on glass slides (Superfrost® Plus, VWR) and stored at −20°C. For immunostaining, cryosections on glass slides were air-dried for one hour and then washed three times for 5 min with PBS. Afterwards, the sections were stained following the same protocol as the cells (see above).

### 2.3 Image analysis

Cell cultures were first immunostained and imaged for the proteins of interest. When expression levels were compared between conditions the same acquisition settings were used for all images. Then, at least 30 cells were chosen at random for each condition to analyze the fluorescence intensity of the whole cell using the Fiji software [59]. All data were analyzed using unpaired *t*-tests with Welch’s correction using GraphPad Prism software. Each experiment was repeated three times. The results are presented as mean + standard deviation. Significance values were represented as: ** = *P* < 0.01, *** = *P* < 0.001.

## 3. Results

Integrins are ubiquitously expressed throughout the retina, mediating a variety of functions. For instance, on RPE cells, they mediate the interaction with Bruch’s membrane and the phagocytosis of shed POS [1,3,5]. In this study we analyzed and compared the expression of integrin subunits in different RPE cells using immunostaining. We assessed the integrin expression *in situ* using rat eye cryosections, in primary rat RPE cells [55], and in the spontaneously immortalized human RPE cell line ARPE-19 [41] using differentiated and undifferentiated cells [58].

### 3.1 Integrin expression in RPE cells *in situ*

As a first step, we assessed the expression of integrins *in situ*. To this end, we used cryosections of adult SD rat eyes and performed immunostaining targeting integrin subunits. In addition to the integrins already assessed in our previous studies, α1, α3, α5 and αV as well as β1 and β3 [60], we expanded the pallet of integrins evaluated by α6, α8, and β4. As discussed previously [60], the α1 integrin subunit was expressed by a variety of cells within the retina. Expression of α1 was visible in the RPE cells (visualized by red fluorescence [65 kDa RPE specific protein (RPE65); 61,62]), the retinal ganglion cells (RGCs), cell bodies and processes throughout the length of the retina (Figure 1, α1). The α3 integrin subunit was visible in the RPE cells and cell processes in the inner plexiform layer (IPL) and the ganglion cell layer (GCL) (Figure 1, α3). α5 integrins were expressed in the choroid, RPE cells, and throughout the whole length of the retina (Figure 1, α5). No expression of the α6 integrin subunit was detectable in the retina. However, this integrin subunit was visible within the sclera and the choroid and to some extent in RPE cells (Figure 1, α6). The α8 integrin subunit was clearly detectable in the inner nuclear layer (INL) and the RGC layer with some immunoreactivity visible in the RPE cell layer (Figure 1, α8). RPE cells, cells in the INL and IPL, and RGCs expressed the αV integrin subunit (Figure 1, αV). β1 integrin subunit expression was obvious in the RPE layer, the INL and the RGC layer (Figure 1, β1). Immunoreactivity of the β3 integrin subunit was visible in the RPE cells and cell processes in the IPL and GCL (Figure 1, β3). The β4 integrin subunit was found in the RPE cells, the INL and the RGC layer (Figure 1, β4). In summary, RPE cells *in situ* expressed all the integrin subunits tested.

**Figure 1:**
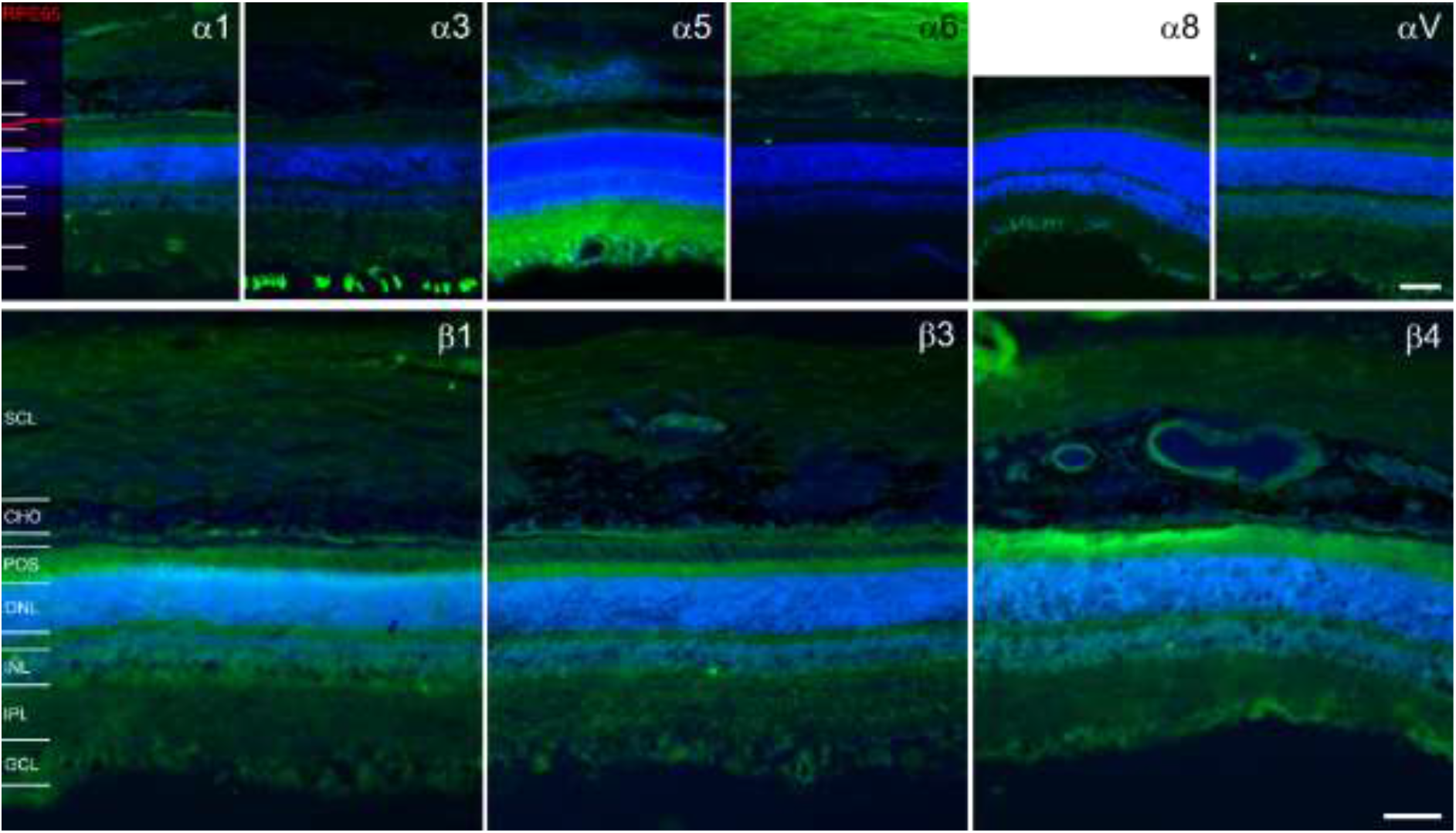
Expression of integrin subunits in the rat retina. (**α1**) Immunostaining for the α1 integrin subunit (green) revealed its presence in the RPE cells and the RGCs as well as cell processes throughout the length of the retina. RPE cells are labeled with RPE65 (red). (**α3**) The α3 integrin subunit (green) was expressed in the RPE cells, cell processes in the IPL and in RGC cell bodies. (**α5**) Expression of the α5 integrin subunit (green) was illuminated in the choroid, the RPE cells, the INL and IPL and the RGC layer. (**α6**) The α6 integrin subunit (green) was expressed by the RPE cells, and cells in the choroid and the sclera. (**α8**) Immunostaining showed expression of the α8 integrin subunit (green) in the INL and the RGC layer. Some immunoreactivity was also detectable in the RPE cell layer. (**αV**) The αV integrin subunit (green) was expressed by the RPE cells, cells in the INL and IPL, and the RGCs. (**β1**) The β1 integrin subunit (green) was visualized in the RPE cells, the INL and the RGC layer. (**β3**) The β3 integrin subunit (green) was expressed by the RPE cells and cell processes in the IPL. (**β4**) RPE cells, cells in the INL and the RGC layer expressed the β4 integrin subunit (green). Cell nuclei are labelled with DAPI (blue) throughout. SCL = sclera, CHO = choroid, POS = photoreceptor outer segments, ONL = outer nuclear layer, INL = inner nuclear layer, IPL = inner plexiform layer, GCL = ganglion cell layer. Scale bars = 50 μm.

### 3.2 Integrin expression in primary adult rat RPE cells

Next, we cultured adult rat RPE cells as described previously [55] and assessed their integrin expression. We used SD rat-derived cells to avoid autofluorescence from the pigments as these rats are albino. We cultured the cells for two weeks before fixation and immunostaining. The RPE cells formed a cobble-stone like monolayer in culture, a hallmark for RPE cells (Figure 2, A), and expressed the RPE cell markers cytokeratin 18 (Figure 2, B) and zonula occludens-1 (ZO-1) (Figure 2, C) confirming their lineage [62–64].

**Figure 2:**
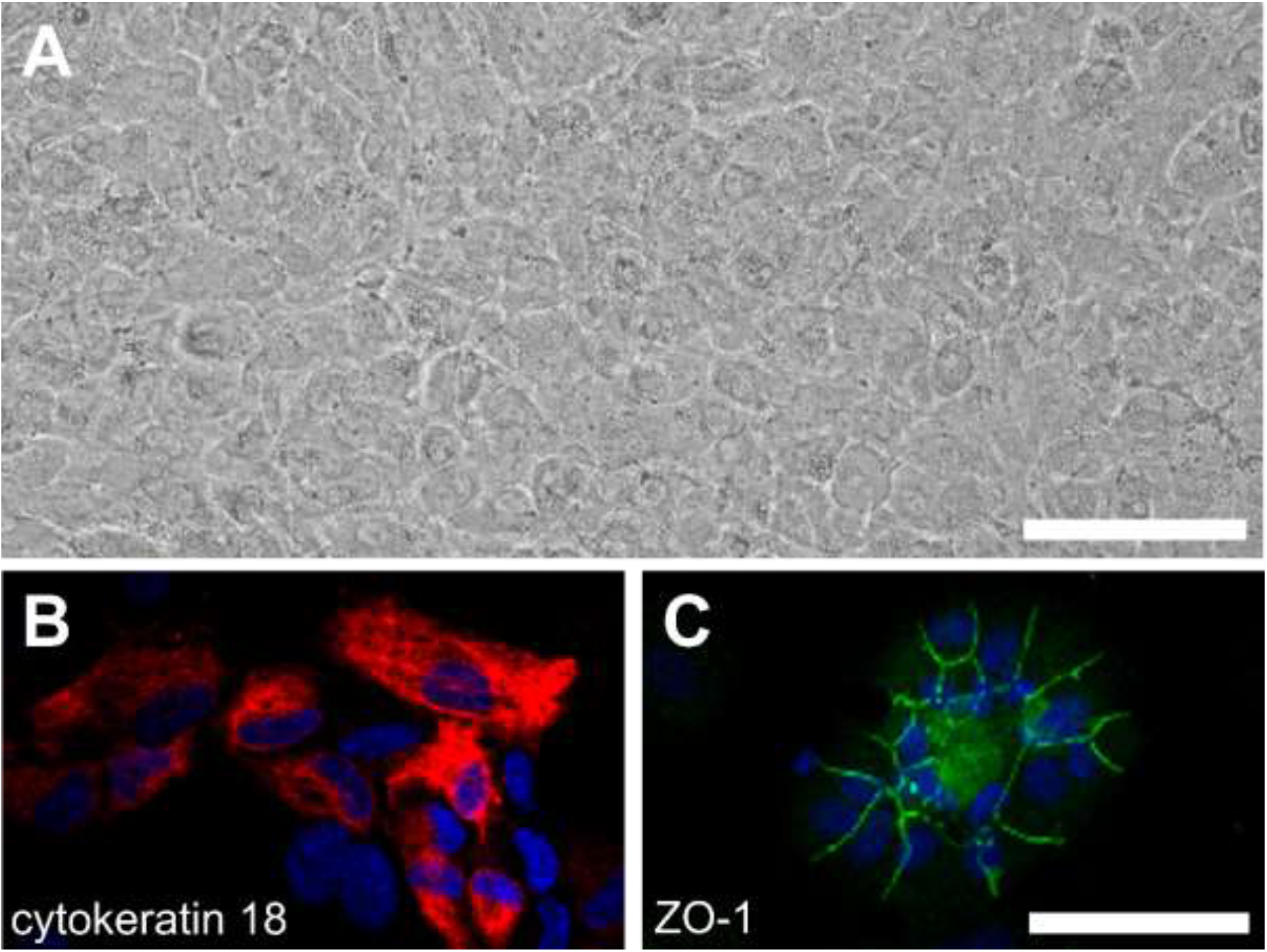
Cultured primary adult rat RPE cells. **(A)** SD rat-derived RPE cells formed a non-pigmented monolayer in culture. **(B and C)** The cells expressed the RPE cell markers cytokeratin 18 (red) and ZO-1 (green). Cell nuclei were labelled with DAPI (blue). Scale bar = 100 μm.

We performed immunostaining on SD rat-derived RPE cells to visualize integrin subunits expressed by the cells. The immunostaining revealed varying degrees of expression of the α1, α2, α3, α4, α5 and αV, β1, β4 and β5 integrin subunits as well as of αVβ3 and αVβ5 integrins (Figure 3). Especially the immunostaining against the α8 and β3 integrin subunit showed a lot of background signal. On the other hand, the α3, α5 and αV as well as the β1 and β4 integrin subunits were localized to large focal adhesions (intense staining around edges of the cells, marked by arrows) in the cultured RPE cells (Figure 3), confirming their role in the adhesion of the cells.

**Figure 3:**
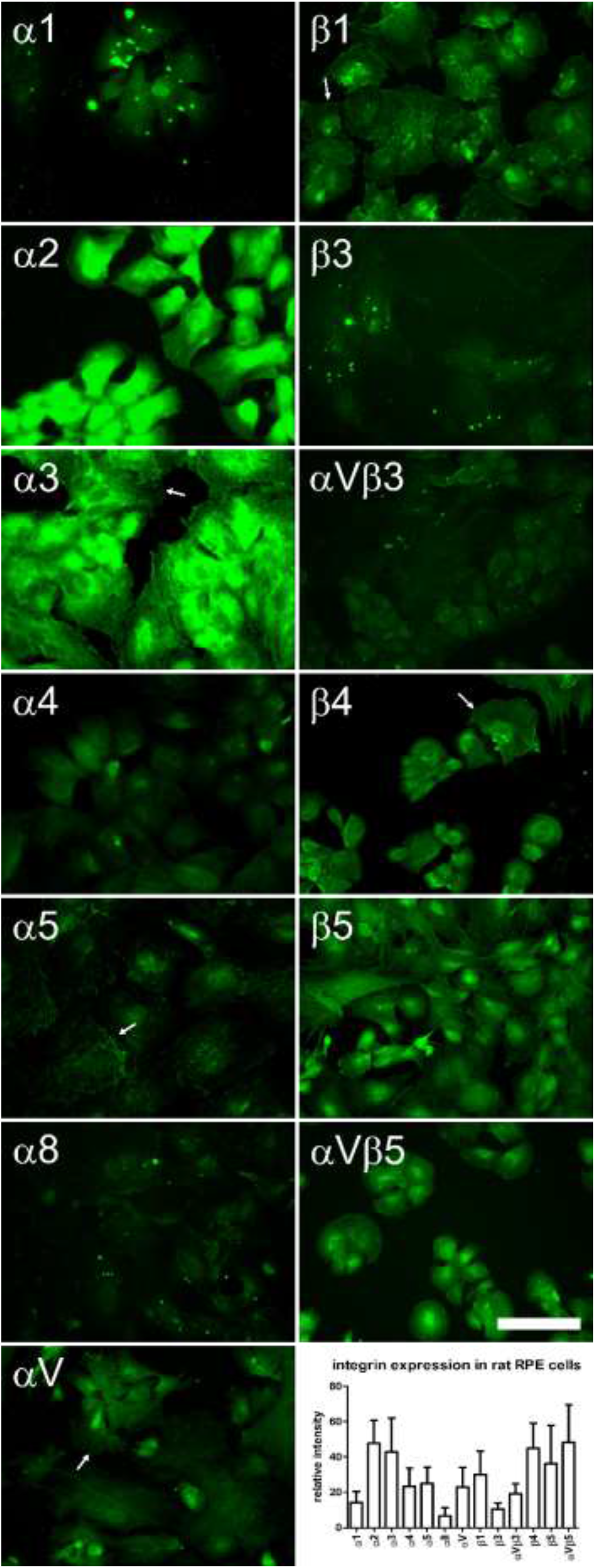
Integrin expression in cultured primary rat RPE cells. The cultured RPE cells expressed α1, α2, α3, α4, α5 and αV, β1, β4 and β5 integrin subunits as well as αVβ3 and αVβ5 integrins. The signals for the α8 and β3 integrin subunits were very weak and might be due to unspecific binding of the antibodies. However, the labelling of the αVβ3 integrin appeared specific as it was targeted to the plasma membrane. The α3, α5 and αV as well as the β1 and β4 integrin subunits were found in large focal adhesions (arrows). Scale bar = 100 μm.

### 3.3 Integrin expression in ARPE-19 cells

After establishing the integrin expression in primary rat RPE cells we used immunostaining to visualize the presence of integrins in the widely used human RPE model cell line ARPE-19 [41]. The cells resembled epithelial cells in culture 24 h after plating (Figure 4, A), and they formed a monolayer after several days in culture (Figure 4, B). Culturing ARPE-19 cells in DMEM^+^ [58] enabled the cells to redifferentiate and to form a tight monolayer after several months in culture (Figure 4, C). The cells in DMEM^+^ started to get pigmented after ~five months and continued to do so until the whole monolayer was pigmented after ~seven months (Figure 4, D).

**Figure 4:**
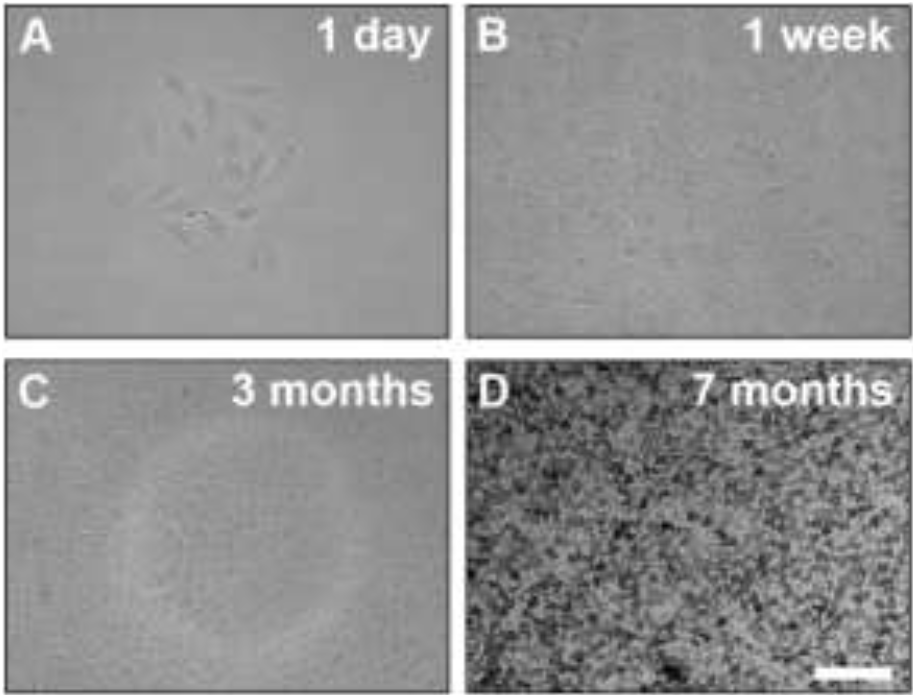
Culture of ARPE-19 cells. **(A)** ARPE-19 cells resembled epithelial cells in culture 24 h after plating. **(B)** The cells formed a disorganized monolayer after one week in culture. **(C)** ARPE-19 cells cultured in DMEM^+^ [58] re-differentiated and formed a tight monolayer after several months. **(D)**The monolayer started to get pigmented after ~five months in culture, and all cells were pigmented after ~seven months. Scale bar = 100 μm.

Next, we used immunocytochemistry to assess the expression of RPE-specific markers in ARPE-19 cells cultured in DMEM^+^ (Figure 5). We used undifferentiated (ARPE-19 cells cultured for two weeks) and differentiated ARPE-19 cells that were cultured for ~five months. Low levels of bestrophin [BEST; 65,66], cellular retinaldehyde binding protein [CRALBP; 61] and RPE65 were visible in the undifferentiated ARPE-19 cells (Figure 5). However, the cells expressed cytokeratin 18 and microphthalmia-associated transcription factor [MiTF; 67,68] (Figure 5). Moreover, immunostaining for the Na^+^/K^+^-ATPase [69] revealed strong immunoreactivity in the perinuclear region of the undifferentiated ARPE-19 cells (Figure 5). The tight junction marker ZO-1 was localized to the cell-cell-contacts but immunostaining exposed a non-organized staining pattern (Figure 5). On the other hand, the differentiated ARPE-19 cells expressed significantly higher levels (unpaired *t*-tests with Welch’s correction: *** = *P* < 0.001) of all the RPE-specific markers tested (Figure 5), resembling native RPE cells [55].

**Figure 5:**
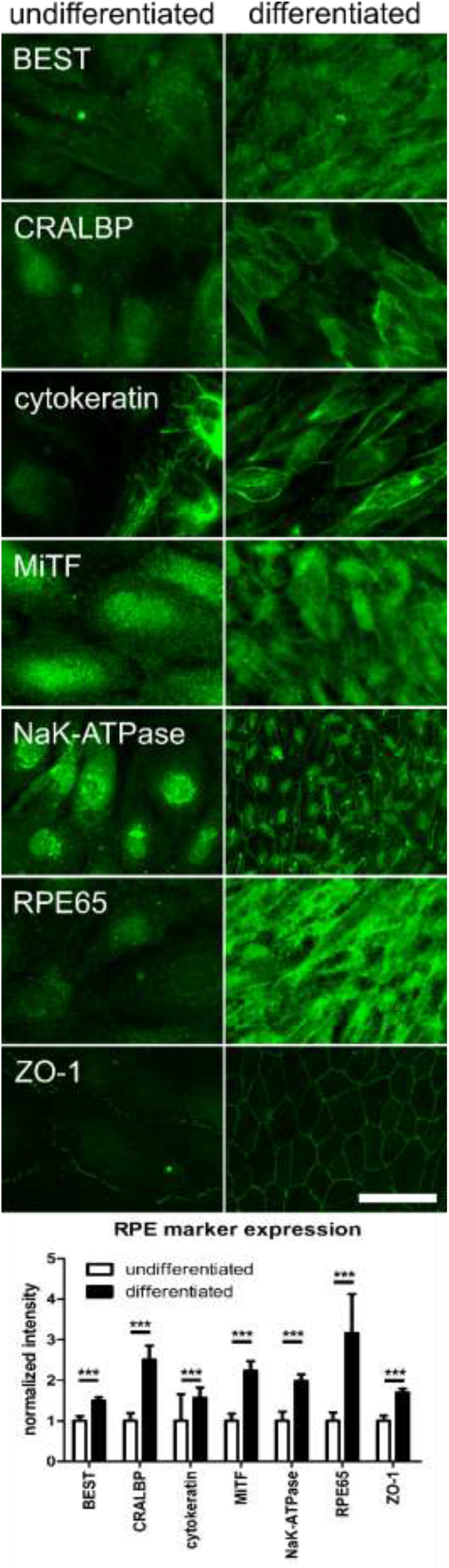
Expression of RPE-specific markers in ARPE-19 cells. Undifferentiated (cells cultured for two weeks) and differentiated ARPE-19 (cells cultured for ~five months) were used for immunostaining to assess the expression of RPE-specific markers. The undifferentiated ARPE-19 cells expressed low levels of BEST, CRALBP and RPE65. However, they expressed cytokeratin 18, MiTF, the Na^+^/K^+^-ATPase and the tight junction marker ZO-1. The differentiated ARPE-19 cells expressed significantly higher levels of all the RPE-specific markers tested (unpaired *t*-tests with Welch’s correction: *** = *P* < 0.001). Scale bar = 50 μm.

After confirming the expression of RPE-specific markers in ARPE-19 cells we next assessed the integrin subunit expression in these cells (Figure 6). Again, we used undifferentiated and differentiated cells (see above). Although only low fluorescence signals with high background were detectable for the α1, α5 and α8 integrin subunits, the undifferentiated ARPE-19 cells expressed the α2, α3, α4 and the αV integrin subunits (Figure 6). In contrast, the differentiated ARPE-19 cells expressed all α integrin subunits tested (Figure 6). Again, only low levels of the α1 integrin subunit were found. However, the differentiated ARPE-19 cells showed an expression of the α5 and the α8 integrin subunits (Figure 6). In addition to α integrin subunits, the undifferentiated ARPE-19 cells expressed the β1 integrin subunit and the αVβ3 integrin on their surface although the staining for the β3 integrin subunit alone was weaker (Figure 6). The undifferentiated ARPE-19 cells expressed the β4 integrin subunit which displayed strong staining in the nucleus (Figure 6). Moreover, the cells expressed the β5 and the αVβ5 integrin on their cell surface (Figure 6). On the other hand, the differentiated ARPE-19 cells expressed the β1, β3, β4 and β5 integrin subunits as well as the αVβ3 and αVβ5 integrins on their surface (Figure 6). With the exception of the β3 integrin subunit, the differentiated ARPE-19 cells expressed significantly higher levels (unpaired *t*-tests with Welch’s correction: ** = *P* < 0.01 and *** = *P* < 0.001) of all integrins assessed (Figure 6).

**Figure 6:**
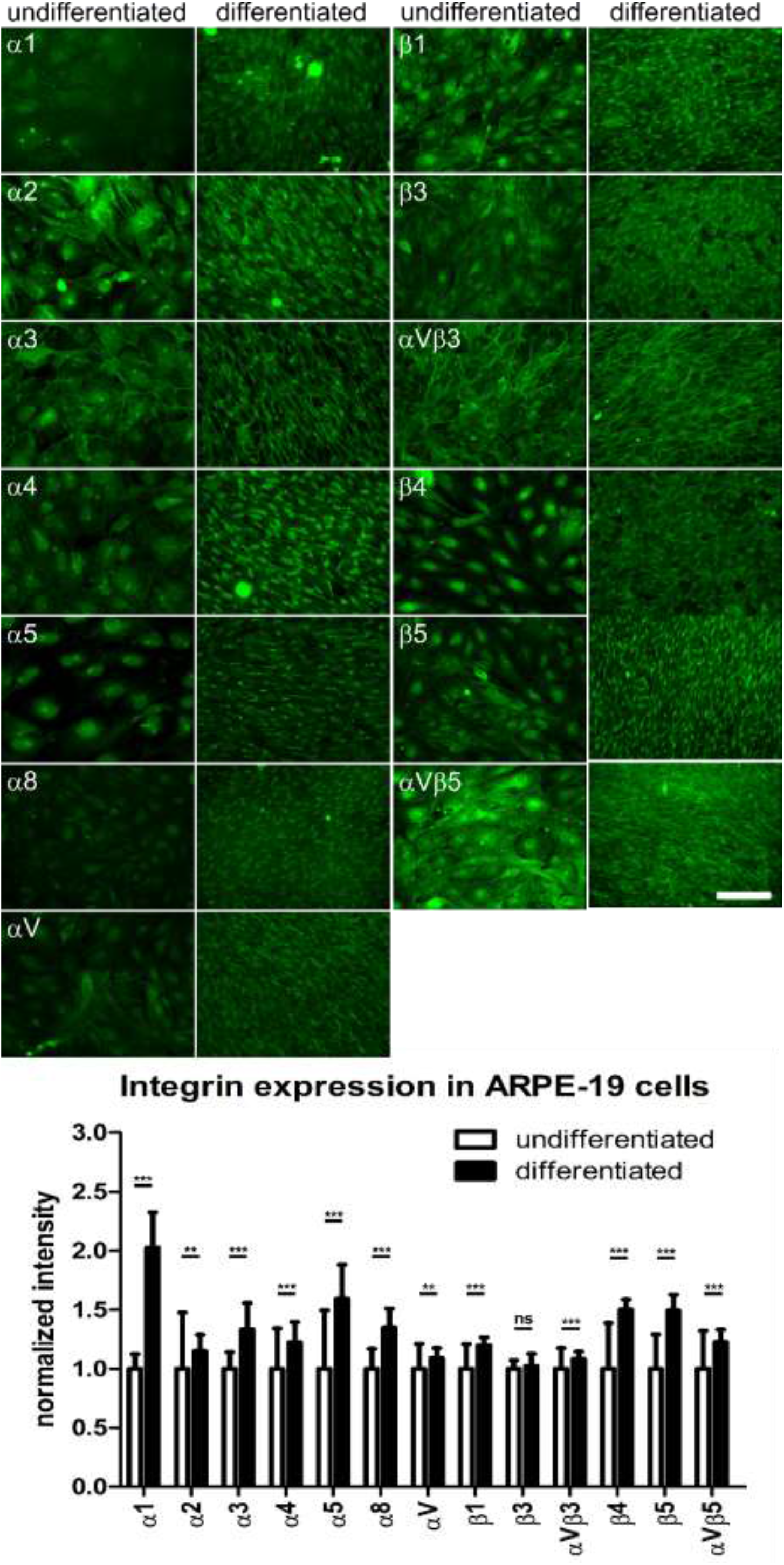
Expression of integrin subunits in ARPE-19 cells. We performed immunostaining on undifferentiated and differentiated ARPE-19 cells to assess the presence of integrin subunits. Undifferentiated ARPE-19 cells expressed the α1, α2, α3, α4, α5, α8 and αV integrin subunits. The immunoreactivity for the α1, the α5 and the α8 integrin subunit was very faint. Also the differentiated ARPE-19 cells expressed low levels of the α1 integrin subunit. However, they expressed the α2, α3, α4 and αV integrin subunits as well as α5 and α8 integrin subunits. The α5 subunit displayed a staining pattern different to the other integrin subunits tested and revealed long stretches of the protein on the cell surface. Moreover, the α8 integrin subunit immunoreactivity was mostly visible in the nucleus. The undifferentiated ARPE-19 cells expressed the β1, β3, β4 and β5 integrin subunits as well as the αVβ3 and αVβ5 integrins. However, the fluorescence signal for the β3 integrin subunit was very weak, and the β4 integrin subunit was mostly visualized in the cell nuclei. The differentiated ARPE-19 cells expressed the β1, β3, β4 and β5 integrin subunits as well as the αVβ3 and αVβ5 integrins on their surface. With the exception of the β3 integrin subunit, the differentiated ARPE-19 cells expressed significantly higher levels (unpaired *t*-tests with Welch’s correction: ** = *P* < 0.01 and *** = *P* < 0.001) of all integrins assessed. Scale bar = 100 μm.

## 4. Discussion

In this study we investigated the expression of integrin subunits in the retina *in situ*, in primary adult rat RPE cells and in the human RPE cell line ARPE-19 using immunostaining. Moreover, we evaluated the impact of long-term culturing of ARPE-19 cells on the expression of RPE markers as well as integrin subunits.

The immunostaining results of the integrin expression in the retina are in line with previously published data. The α1 integrin subunit has been located in RGCs [70] and the RPE layer [4,60,71]. Also the α3 integrin subunit has been shown to be expressed throughout the retina [60]. Main points of expression are the RPE cells [4,71,72] and Müller glia, synaptic layers as well as amacrine and some cone cells [73]. Therefore, the staining in the RGC layer in Figure 1 might reflect the Müller cell endfeet as well as displaced amacrine cells. It was demonstrated that the α5 integrin subunit is expressed by the RPE cells [4,74–76]. Furthermore, α5 integrin subunit staining has been demonstrated in the outer plexiform layer (OPL), outer nuclear layer (ONL) and INL, as well as in the RGC layer similar to our results [60,73]. The α6 integrin subunit has only been identified in the developing neural retina [72,77]. Additionally, some studies showed its expression in RPE cells similar to our findings [71,78]. Fitting with our results, expression of the α8 integrin subunit has been localized to RGC axons and cells in the INL [79,80]. In addition, we saw some expression in the ONL as well as the RPE cell layer which has not been reported yet. The αV integrin subunit has also been studied intensively in the retina [60]. It is expressed in RPE cells [4,6,74,81,82] and by cells in the INL and ONL, OPL, the POS layer as well as in RGCs [82], which matches our findings.

Besides the α integrin subunits we localized three β integrin subunits in the rat retina. β1 integrin subunit expression has been demonstrated in RPE cells [4,75,83], RGCs and other neurons in the retina as well as Müller cells [60,80,84] which suggests a similar staining pattern to our results. Also the β3 integrin subunit has been visualized in the retina [60,84] with a weak staining in the nerve fiber layer [72]. In addition, the β3 integrin subunit is expressed by RPE cells [81,85,86] as seen in our immunostaining experiments. The β4 integrin subunit is expressed by the RPE cells and forms a heterodimer with the α6 integrin subunit. The α6β4 integrin links laminin in the ECM to the keratin intermediate filaments of the cytoskeleton [4,87]. Surprisingly, our staining showed a distribution of the β4 integrin subunit throughout the retina, with strong immunoreactivity in cell bodies in the INL and the RGC layer. However, the β4 integrin subunit has only been shown to dimerize with the α6 integrin subunit [11,13]. Nevertheless, our α6 integrin subunit staining did not show any immunoreactivity in the retina. Potentially, this might be due to the use of an antibody raised in rat and used on rat tissue. Hence, either one of the immunostaining reactions yielded unspecific results or the β4 integrin subunit is able to form a heterodimer with an integrin subunit other than α6. Some published data suggest the existence of an αVβ4 integrin [88,89]. However, the staining for the αV integrin subunit alone did not show a distinct staining pattern in the cell bodies in the INL as seen in the staining for the β4 integrin subunit. Thus, the β4 integrin subunit might interact with a yet unknown dimerization partner.

In this study, we used immunostaining to assess the presence of integrin subunits in cultured primary adult rat RPE cells isolated using papain. The cells expressed several α (α1-5, α8 and αV) as well as β (β1, β3-β5) integrin subunits which confirmed previous studies (see above). However, we did not perform gene expression analyses. Several mouse expression profiles have been published already, confirming the integrin expression pattern [90–92].

The human RPE cell line ARPE-19 has been used as a model for RPE cells for several decades [29,37,40,41,86,93]. However, the cells are non-pigmented, and it is difficult to detect essential RPE markers such as RPE65 in the cells [41,46,58,94–97]. Using immunostaining we were only able to clearly visualize cytokeratin 18, MiTF, Na^+^K^+^-ATPase and ZO-1 in undifferentiated ARPE-19 cells. This is in agreement with other studies [58]. However, upon differentiation using long-term culturing in DMEM^+^ (culturing for at least five months) the cells became pigmented and started to express RPE markers BEST, CRALBP and RPE65 (Figure 5) [58]. Ahmado and colleagues reported the pigmentation of ARPE-19 cells in DMEM^+^ after three months [58]. Low serum and the presence of pyruvate induce the pigmentation of the ARPE-19 cells. Additionally, the expression levels for CRALBP, MERTK and RPE65 are highest under pyruvate conditions [58]. It is therefore important to indicate in which state of differentiation the ARPE-19 cells are as each state shows distinct phenotypes and expresses a different battery of genes.

Using immunostaining we identified several α and β integrin subunits in ARPE-19 cells. One has to bear in mind however that the culture medium [76], the culture substrate [98] as well as the passage number [99] influence gene expression. Proulx *et al*. showed that the expression of the α5 integrin subunit declines when cells reach post-confluency levels [75,76]. Similarly, in our immunostaining study the staining for the α5 integrin subunit changed when comparing undifferentiated and differentiated ARPE-19 cells. Even though the overall staining intensity increased, strong immunoreactivity at cell-cell borders that appeared as ‘worm-like’ structures became visible. However, with the exception of the β3 integrin subunit, the expression of the other integrin subunits tested increased during the transition of the ARPE-19 to a pigmented and more native phenotype. This was especially visible for the α1, the β4 and the β5 integrin subunits as well as for the αVβ5 integrin. The α1 and the β4 integrin subunits are involved in the adhesion of the RPE cells to laminin [87,100], an ECM molecule that is produced by RPE cells [55,78]. Additionally, as mentioned, the αVβ5 integrin is essential for RPE cell function, *i.e*. binding of POS and subsequent phagocytosis through CD36 and MERTK [3,6–9]. Our results also fit with gene expression profiles that have been published for ARPE19 cells [101], fetal and adult human RPE cells [102,103] as well as stem cell-derived RPE cells [104,105].

All RPE cells tested expressed integrins which enable them to interact with normal Bruch’s membrane. However, as mentioned above, the composition of the Bruch’s membrane changes during aging and in disease making it more anti-adhesive [14–17]. Moreover, the removal of choroidal new vessels in the treatment of AMD leads to the exposure of deeper layers of Bruch’s membrane [106–109] that do not favor the attachment of RPE cells [26,27,29,30,110]. In addition, in wet AMD an upregulation of the inhibitory glycoprotein tenascin-C (TN-C) occurs on Bruch’s membrane [86,111,112] in addition to changes in proteoglycan composition, which might also render the membrane less adhesive [113]. It has been shown that RPE cells express at least one receptor that normally binds to TN-C, αVβ3 [86]. αVβ3 can be activated through manganese and blocking β3 integrins reduces ARPE-19 binding to and migration on TN-C [86]. Surprisingly, we found the expression of the α8 integrin subunit in RPE cells. The α8β1 integrin is a receptor for TN-C [114,115] and should mediate the adhesion of the RPE cells to TN-C. However, we only detected low levels of this integrin subunit in the RPE cells, similar to gene expression analyses mentioned above. The α8β1 integrin might be activated together with αVβ3 integrin to enable binding to TN-C. Nonetheless, the activation of β1 integrins alone did not increase ARPE-19 adhesion to TN-C [86].

Studies showed that β1 integrins are the most important receptors for the adhesion and migration of RPE cells and that their blocking leads to a failure of RPE cell attachment and migration [116–118]. In addition, blocking α integrin subunits on RPE cells results in less spreading and adhesion of the cells on Bruch’s membrane explants and components [71]. Uncultured RPE cells express low levels of integrins on their surface which can impair the reattachment of the cells to Bruch’s membrane or other surfaces [4]. As seen in our results, upon culturing however, the levels of surface integrins rise and the cells bind better to Bruch’s membrane [4,71].

## Acknowledgments

Funding: This work was supported by grants from the Medical Research Council (G10000864) (JH, TH, JK and JF), Christopher and Dana Reeves Foundation (JF), the Wellcome Trust MRC Cambridge Stem Cell Institute and Fight for Sight (KM), Czech Science Foundation GACR 19-10365S, project “Center of Reconstructive Neuroscience”, CZ.02.1.01/0.0./0.0/15_003/0000419, a Wellcome Trust Principal Fellowship (101896) (JH), the European Union’s Horizon 2020 research and innovation program (Marie Skłodowska-Curie grant agreement 798644-AstroMiRimage) (JH) and a research grant from Science Foundation Ireland (SFI, 16/RC/3948) and co-funded under the European Regional Development Fund and by FutureNeuro industry partners (JH).

## Author Contributions

Conceptualization, JH, KM and JF; Methodology, JH, JK; Formal Analysis, JH; Investigation, JH and TH; Writing – Original Draft Preparation, JH; Writing – Review & Editing, JH, TH, JK, KM and JF; Supervision, JK, KM and JF; Funding Acquisition, JF

